# Actin pool controls the scaling of dynamic actin networks

**DOI:** 10.64898/2026.11.03.763964

**Authors:** Alice Cantat, Christophe Guérin, Nevena Morel, Benoit Vianay, Alex Mogilner, Manuel Théry, Laurent Blanchoin, Alexandra Colin

**Affiliations:** Interdisciplinary Research Institute of Grenoble, Laboratoire de Physiologie Cellulaire & Végétale, CytoMorpho Lab, University of Grenoble-Alpes, CEA, CNRS, INRA, Grenoble, France; Courant Institute of Mathematical Sciences, New York University, New York, United States; Department of Biology, New York University, New York, United States; Université Paris Sciences et Lettres, CEA, ESPCI, Institut Pierre-Gilles de Gennes, CytoMorpho Lab, Paris, France

**Keywords:** Actin, reconstituted systems, scaling, robustness, adaptability, limited pool, depletion

## Abstract

Scaling is a fundamental property of living systems, allowing cells to match the size of their internal components to their volume. The size and shape of cells depend largely on the actin cytoskeleton, whose dynamic self-assembling structures are central to this adaptation. Yet how they self-assemble in a volume-dependent manner remains unresolved. To address this question, we used a biomimetic system consisting of beads propelled by actin polymerization in microwells of different volumes, which allowed us to work with a limited and defined number of components. This approach enabled the first reconstitution, from purified proteins, of dynamic cytoskeletal structures that adapt to the volume of their environment. The scalability of these structures depends on how strongly they deplete monomers from a shared pool, and tuning the disassembly rate is sufficient to shift their scaling regime. Scaling principles also hold for multiple competing structures, with greater competition imposing stronger constraints on the monomer pool. Together, these results establish a mechanistic framework in which a structure’s capacity to adapt is set by how much it draws down shared resources as it grows, a principle that may extend to other intracellular structures that scale with the cells that build them.

## Introduction

Biological systems must continuously adapt their architecture to changes in size while preserving function. At the cellular level, metabolic activity, transport processes, and force production must adapt to changes in cell size in order to maintain function (Cadart & Heald, 2022; Chan & Marshall, 2012; Chen & Levy, 2023; Gu & Oliferenko, 2021; Marshall, 2002, 2015, 2020). Interestingly, intracellular structures do not all obey the same scaling laws. Some organelles exhibit robust proportional scaling with cell size, while others remain largely invariant. The nucleus, for example, scales remarkably linearly with cell volume across a broad range of organisms and cell types. On the other hand, centrioles do not scale with cell size, neither in size nor number (Marshall, 2020). Another well-known observation is Kleiber’s power law that states that metabolism scales to ¾ of body mass (West *et al*, 1997). This diversity in scaling behavior suggests that scaling laws are tied to the functional role of each structure. Therefore, understanding how biological structures scale is necessary to provide insight into the fundamental principles underlying cellular organization.

Among the structures whose scaling is most tightly coupled to function, the cytoskeleton stands out. It governs force generation, intracellular organization, polarity, and division, roles inherently sensitive to cell size and must therefore adapt to cell-size changing events through yet unresolved mechanisms. The mitotic spindle has been extensively studied in this context, and in several systems was shown to scale linearly with cell size before reaching an upper limit for larger cell volumes (Lacroix *et al*, 2018; Good *et al*, 2013; Hazel *et al*, 2013; Wühr *et al*, 2008; Mitchison *et al*, 2015; Reber & Goehring, 2015; Kletter *et al*, 2025; Rieckhoff *et al*, 2020; Farhadifar *et al*, 2015). In the case of actin, it has been shown that the size of the structures adapts differently depending on their function. In fission yeast, actin cables length correlate with cell length (McInally *et al*, 2021), while number of stress fibers correlate with cell area in epithelial tissue (López-Gay *et al*, 2020). During cytokinesis, the number of precursor nodes contributing to the contractile ring scales with cell volume (Sayyad & Pollard, 2022). This variability in scaling laws may reflect that of the functions and dynamics of actin structures. However, the fundamental mechanisms underlying the scaling laws of actin structures remain largely unknown.

Understanding scaling laws ultimately requires understanding how the size of a structure is controlled. The simplest way to impose control on a structure or process is through a feedback loop (El-Samad, 2021; Banerjee & Banerjee, 2025). Several mechanisms have been proposed (Marshall, 2020; Reber & Goehring, 2015; Goehring & Hyman, 2012), but in the context of the cytoskeleton, and actin structures in particular, the limiting pool is the best known feedback (Goehring & Hyman, 2012). In brief, as a structure grows it incorporates subunits from a finite pool, progressively depleting it; because assembly rates depend on the concentration of free subunits, this depletion slows assembly, generating a negative feedback loop capable of controlling structure size. Because actin structures are dynamic and typically undergo continuous assembly and disassembly, or turnover, their size also depends on disassembly rates. This gives rise to a balance-point mechanism for size control, in which a stable size is reached when assembly and disassembly rates match (Banerjee & Banerjee, 2022; McInally *et al*, 2021; Reber & Goehring, 2015). Although disassembly mechanisms differ from one actin structure to another (Mohapatra *et al*, 2016; Rosario *et al*, 2023), these studies converge on a common constraint: at least one of the feedbacks, on assembly or on disassembly, must be length-dependent for robust size control to be achieved. This requirement can be naturally extended to scaling: at least one feedback must be cell-size dependent for scaling to emerge. These models have also revealed the limits to the limiting pool mechanism: showing that when structures compete, the limiting pool feedback alone cannot achieve robust size control (Mohapatra *et al*, 2017). Yet competing actin structures do co-exist in the cell, and there is evidence for a shared subunit pool (Antkowiak *et al*, 2019; Burke *et al*, 2014; Suarez *et al*, 2015; Suarez & Kovar, 2016; Kadzik *et al*, 2020). A recent model indicates that disassembly plays a crucial role in size control within such competitive environments (Momin *et al*, 2026). Despite this progress, the mechanisms controlling the size of actin structures and their consequences for actin scaling laws remain experimentally unresolved.

Resolving these mechanisms *in vivo*, however, poses substantial challenges. As noted above, multiple actin structures co-exist and draw on a common subunit source, yet display distinct scaling laws and size control mechanisms. Studying one structure without perturbing the others or without the measurement being confounded by them is inherently difficult. To address cellular processes free of this surrounding complexity, a number of reconstituted assays have been developed, notably for cytoskeletal questions (Cantat & Colin, 2025). Several studies pertaining to the actin cytoskeleton have used ranges of environment sizes, notably cell-sized droplets, as a mean to study the effect of geometrical confinement on several processes such a contracting waves for example (Krishna *et al*, 2024; Malik-Garbi *et al*, 2019; Miyazaki *et al*, 2015; Sakamoto *et al*, 2020). Very few, however, examined confinement through the lens of the limiting-pool feedback (Colin *et al*, 2023; Guérin *et al*, 2025), nor addressed the scaling of actin structures under this premise.

Here, we have reconstituted an actin system that can adapt to the size of its environment in the simplest possible way, using purified proteins. We placed a constant source of actin nucleation, a bead, inside microfabricated microwells that act as cell-sized compartments of defined volume. This gave us a simple, fully controlled setting in which to observe scaling emerge. Without disassembly, growing branched networks scale linearly with the size of their compartment. Adding ADF/cofilin and Cyclase-Associated-Protein (CAP) to drive disassembly and recycling shifts this to a sublinear scaling regime.

We further show that simply tuning how fast the actin network is disassembled can change the scaling regime. Most strikingly, when structures compete for a shared, limited pool of subunits, this competition alone can set their size. Together, these results reveal general principles that govern the size control of competing dynamic actin architectures.

## Results

### In the absence of turnover, actin structures scale linearly with microwell volume

To investigate how scaling affects branched actin networks, we first reconstituted their assembly in the absence of disassembly, comparing microwells of different sizes to a bulk condition providing effectively unlimited resources (**Figure 1A**). Contrasting these two situations let us isolate the effect of a limited monomer pool on network growth.

**Figure 1.**
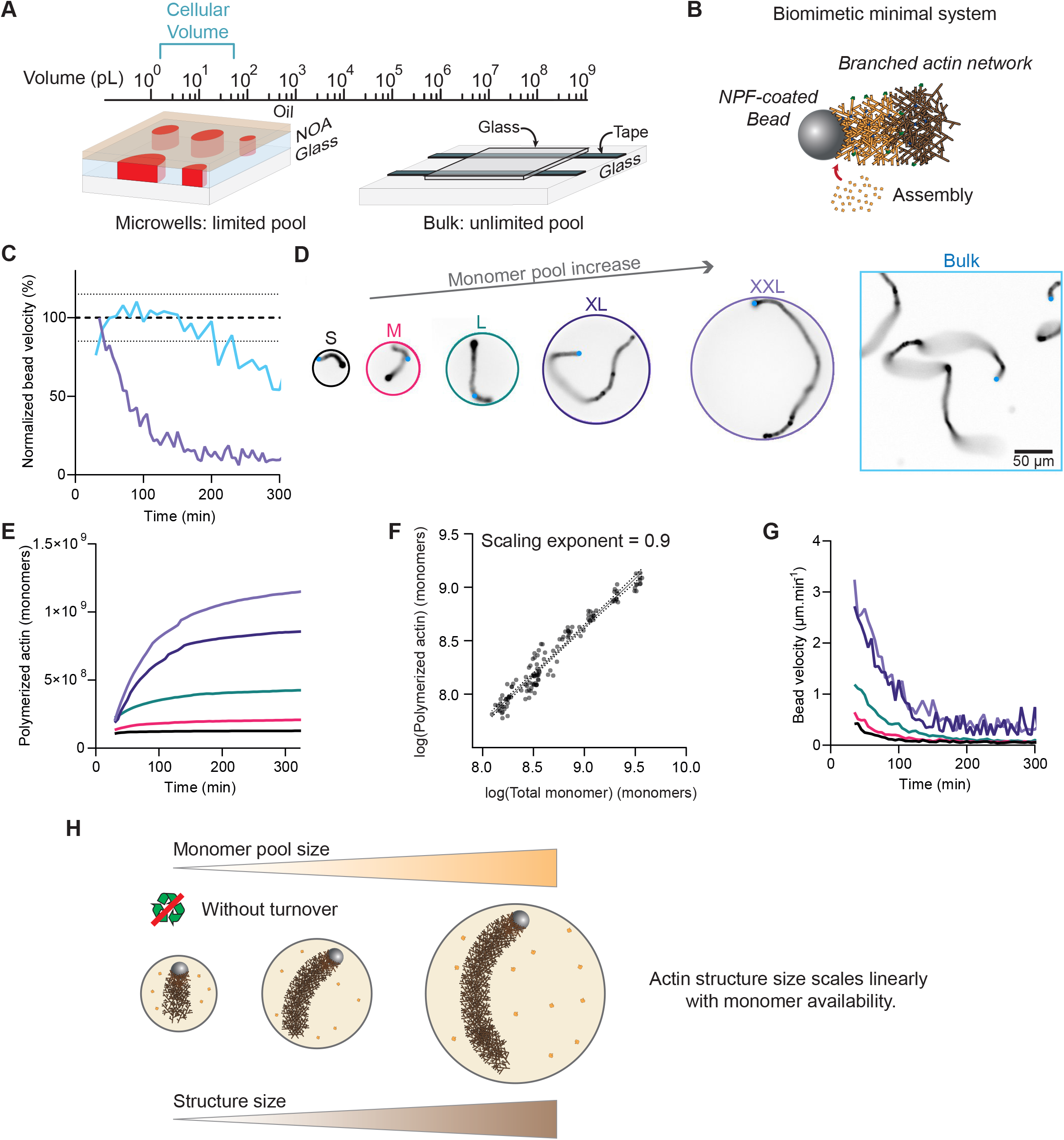
Actin structures growing in conditions without turnover scale linearly with increasing microwell sizes. A) Scheme of the microfabricated microwells and bulk environments. B) Scheme of the bead motility assay: 4 µm polystyrene bead coated with Nucleation-Promoting-Factor (NPF) triggers actin branched network assembly from a mix of actin, profilin, Arp2/3 complex and capping protein. C) Normalized bead velocity to the bulk average overtime. D) Snapshots of actin structures in conditions without turnover in the 5 sizes of microwells (diameter = 50, 75, 100, 150, 200 µm), at 270 min, in grey the actin channel, in blue the bead. Scale bar: 50 µm. E) Quantity of polymerized actin monomers in comet tails overtime for each microwell size. F) Log/Log plot of polymerized actin as a function of total actin monomers per microwell, taken at 270 min. Curve slope is 0.90, R^2^=0.95. G) Bead velocity overtime, for each microwell size. H) Scheme summarizing the effects of microwell size on the size of assembled structures: initial pool size proportionally determines structure dynamics, and final size upon near complete monomer depletion. *Replicates:* S: N=2, n=42, M: N=46, n=2, L: N=3, n=21, XL: N=1, n=11, XXL: N=1, n=12, Bulk: N=1, n=30. *Biochemical conditions*: 6 µM actin, 12 µM profilin, 60 nM capping protein, 90 nM Arp2/3 complex.

We adapted our previous set-up (Colin *et al*, 2023; Vianay *et al*, 2025) to generate microwells spanning a wide range of volumes, from 70 to 590 pL (**Figure 1A, Figure S1A-C**). Each microwell contained a single bead that served as a nucleation source, driving network growth through the bead motility assay, a well-established minimal system for reconstituting branched actin structures (**Figure 1B**; Colin *et al*, 2023; Guérin *et al*, 2025). A much larger bulk chamber provided the reference condition, in which resources are neither limiting nor depleted. Because the available monomer pool scales with microwell volume, this design let us directly test how network size responds to resource availability and quantify how strongly growth is constrained by the limited pool. In this assay, bead velocity reports on the actin assembly rate, which itself depends on actin concentration (Colin *et al*, 2023). We therefore defined an environment as non-depleted when bead velocity remained constant over time, and as depleted when bead velocity decreased over time (**Figure 1C**). Using this readout, we followed branched actin networks in microwells of different volumes (**Figure 1D, Movie 1**). We measured the fluorescence intensity of actin in the comet tails over time. Using a calibration curve based on precise microwell-volume measurements and a defined quantity of labelled actin (Methods, **Figure S1A-E** (Vianay *et al*, 2025)), we converted this signal into the number of actin monomers per comet tail as a function of time and microwell volume (**Figure 1E**). Across all volumes, comet growth followed the same characteristic behavior: a rapid initial phase that progressively slowed until reaching a plateau (**Figure 1E**). Plotting the quantity of polymerized actin at 270 min (in the stalled phase) against the initial number of monomers available in each microwell revealed a linear relationship, with a scaling exponent of 0.9 (**Figure 1F**). In parallel, bead velocity decreased over time in microwells of all sizes (**Figure 1G**). Together, these observations show that the monomer pool is progressively depleted and that network growth is ultimately set by the amount of monomers initially available.

Overall, these data show that the limiting-pool feedback tightly controls comet size in the absence of turnover. In small microwells, depletion sets in quickly, producing short comets that stall early; in larger microwells, growth continues longer before stalling, yielding longer comets (**Figure 1H**). However, the functions of actin structures rely on their intrinsic dynamics. In our system, adding the disassembly factor ADF/cofilin together with the recycling factor cyclase-associated protein (CAP) is sufficient to reconstitute stable, dynamic structures whose turnover matches that of lamellipodia *in vivo* (Colin *et al*, 2023). We therefore focused on the scaling behavior of actin structures once they become dynamic.

### In presence of fast turnover, actin structures do not scale linearly with microwell volume

We next placed beads in microwells spanning the same range of size as before, but now in the presence of ADF/cofilin and CAP. These conditions produced highly dynamic structures (**Figure 2A, Movie 2**) that were shorter than those formed without turnover. Both comet area and bead velocity remained constant over time, indicating that the structures reached a dynamic steady state that lasted for several hours (**Figure S2A-C**). We therefore defined a measurement window between 90 and 150 min and averaged the measurements for each comet within it. Plotting mean comet area against microwell volume confirmed that comets are shorter when they are dynamic (**Figure 2B**). More strikingly, the log–log plot revealed a qualitative difference between the two regimes: with turnover, structure size scales sublinearly with microwell volume (slope of 0.4), whereas without turnover it scales linearly (slope of 1, **Figure 2B, inset**).

**Figure 2.**
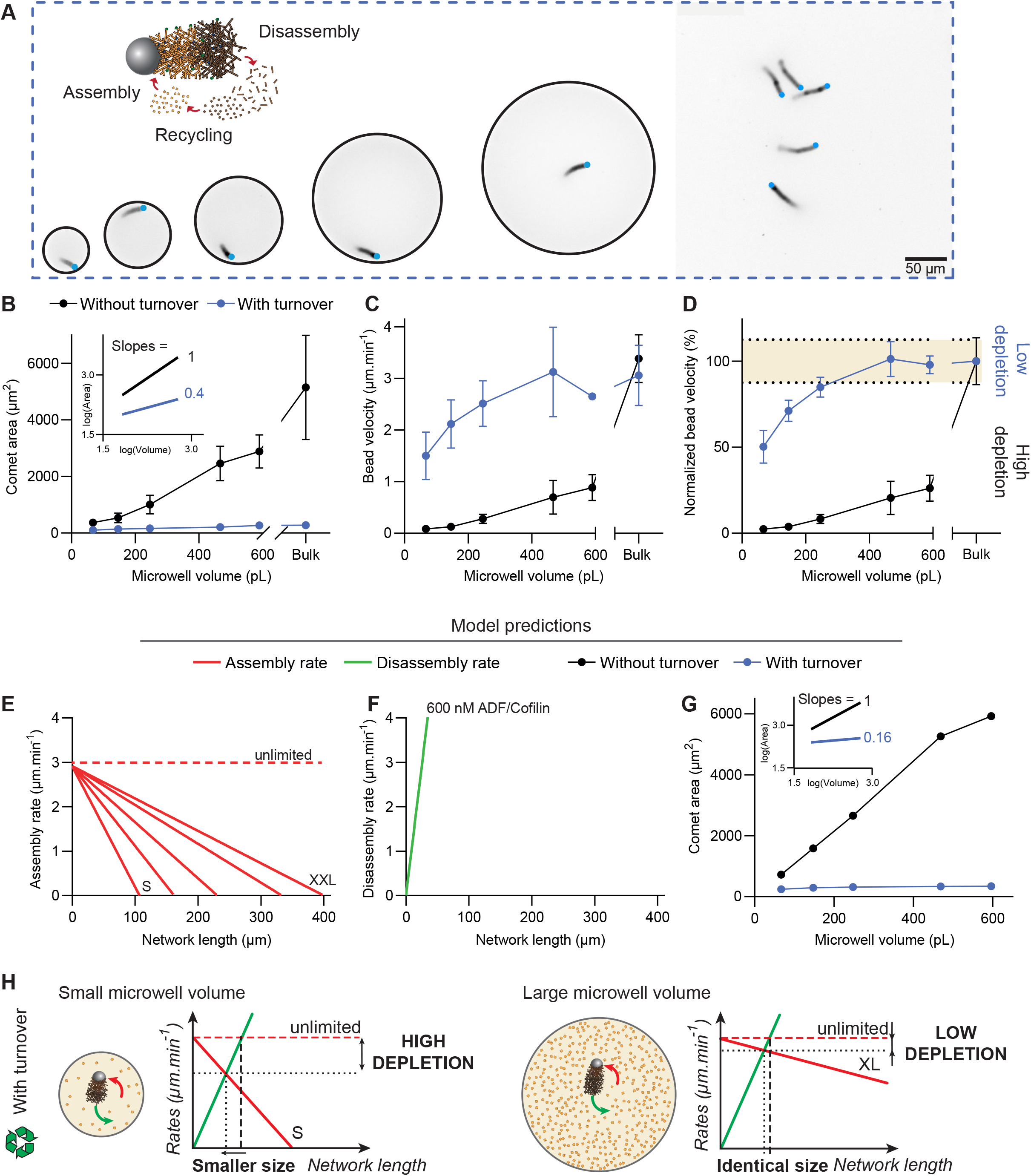
In conditions with turnover, actin structures do not deplete significantly the available pool of monomers in large microwells, leading to sublinear scaling behavior. A) Snapshot of actin structures in turnover in 5 microwell sizes and bulk at 3 hours; in grey is actin, in blue the bead. Scale bar: 50 µm. B) Average comet area between 90 and 150 min for each microwell size and bulk, in conditions without and with turnover. Inset: corresponding log/log plot showing the scaling exponents. C) Average bead velocity between 90 and 150 min, for each microwell size and bulk in conditions without and with turnover. D) Bead velocity normalized to the value in bulk. Colored region: ± 12.5 % around the reference value (100%). E) Model prediction of the assembly rate as a function of network length, for each microwell size and unlimited environment. F) Model prediction of the disassembly rate as a function of network length for turnover conditions at 600 nM ADF/Cofilin and 750 nM CAP. G) Model prediction of the comet area for each microwell size and bulk, in conditions without or with turnover. Inset: corresponding log/log plot and scaling exponents. H) Scheme of the effects of microwell size and turnover conditions on depletion effects: small environments are depleted to a significant extent resulting in smaller sizes and slower assembly. Structures in large environment do not significantly deplete the monomer pool. Color code: Assembly rate model in red, disassembly rate model in green. *Replicates:* S: N=4, n=76, M: N=4, n=80, L: N=4, n=83, XL: N=3, n=14, XXL: N=3, n=12, Bulk: N=3, n=413. *Biochemical conditions*: 6 µM actin, 12 µM profilin, 60 nM capping protein, 90 nM Arp2/3 complex, 600 nM ADF/cofilin, 750 nM CAP.

To understand this sublinear scaling, we asked how the size of a dynamic structure is set. Because structure size reflects the balance between assembly and disassembly rates, we used bead velocity as a readout of the assembly rate in our biomimetic assay (**Figure 1** (Colin *et al*, 2023)), which also reports on how strongly the monomer pool is depleted in microwells of different volumes. With turnover, bead velocity was both higher and closer to the unlimited-environment value than without turnover (**Figure 2C**). To compare depletion across conditions, we normalized each measurement to the unlimited-environment control (**Figure 2D**) and defined a ±12.5% window around the control value, within which velocity is considered comparable to the unlimited case and the monomer pool effectively non-depleted. By this criterion, the larger microwells with turnover (volume >220 pL) showed no detectable depletion. This indicates that the sublinear scaling regime coincides with an absence of monomer depletion in larger environments.

We built a simple kinetic model to explain these observations. In the model, the size of the comet tail is set by the balance between assembly and disassembly (**Methods**). Assembly is proportional to the concentration of available monomers which is equal to the total amount of actin in the microwell minus amount of actin polymerized in the tail, divided by the volume of the microwell. Because the monomer pool is limited, the model predicts that, without turnover, the tail area increases and the bead velocity decreases with time, because gradually more actin is polymerized and fewer monomers are available (**Figure S2D-E**). This limited-pool effect means that the assembly rate falls linearly as the structure grows (**Figure 2E**). Disassembly, by contrast, increases linearly with structure size, with a rate factor γ that depends on the concentrations of ADF/cofilin and CAP (**Methods**; (Manhart *et al*, 2019)). We estimated γ from bulk measurements at steady state (**Methods**). To find the steady-state size of the comet tail, we compared the disassembly rate with the assembly rate as a function of structure size (**Figure 2E-F**). The steady-state size is the point where the two curves intersect, which gives a theoretical estimate of comet size both without and with turnover. Using this approach, the model predicts comet size as a function of microwell volume as well as respective scaling between the volume and the comet size (**Figure 2G**). The close agreement between these predictions and our experimental data indicates that the model can account for what we observe.

The model also explains the sublinear scaling regime observed in the presence of fast turnover. In small microwells, greater depletion lowers the assembly rate, so the structure settles at a smaller size than it would in an unlimited environment (**Figure 2H**). In large microwells, depletion is too insignificant to matter before disassembly balances assembly, so the steady-state size is nearly identical to the unlimited case (**Figure 2H**). When monomer consumption is low, the network reaches a size close to that of a network growing with unlimited resources, and its size no longer depends on compartment volume. This is the sublinear scaling behavior we observe at large volumes. Taken together, the model predicts that the size of the structure is set by how strongly it depletes the monomer pool. If this is correct, varying the extent of depletion should shift the scaling regime from linear, when depletion is high, to sublinear, when depletion is low. To test this, we varied the disassembly and recycling rates by tuning ADF/cofilin and CAP concentrations, sampling a broad range of conditions that affect monomer depletion to different degrees.

### Scalability of dynamic structures depends on their turnover rate

To test how depletion of monomers controls scaling, we varied the concentrations of ADF/cofilin and CAP from 50 nM to 800 nM, spanning low to high turnover rates (**Figure 3A, Figure S3A, Movies 3 and 4**). For each condition, we measured comet area across the full range of microwell volumes (**Figure 3B**). Comets were consistently longer when turnover was low than when it was high. Plotting comet area against microwell volume on a log–log scale revealed that these conditions did not merely shift comet size but changed the scaling behavior itself (**Figure 3B, inset**). To make this explicit, we extracted the scaling exponent for each condition and plotted it against ADF/cofilin concentration, which showed a progressive loss of scaling as the disassembly rate increased (**Figure 3C**). Without turnover, or at low turnover rates (below 200 nM ADF/cofilin), structures scaled linearly with microwell volume (exponent near 1). At higher turnover rates, the exponent dropped below 0.5, indicating that structures no longer scaled linearly with volume. To link this loss of scaling to monomer depletion, we plotted the normalized bead velocity for each turnover rate across microwell volumes (**Figure 3D**). Conditions that showed a sublinear scaling regime (above 400 nM ADF/cofilin) showed very little monomer depletion at large volumes, whereas conditions that scaled linearly (below 400 nM ADF/cofilin) showed strong depletion. Scaling therefore tracks depletion directly: structures scale linearly with the environment volume when the pool is depleted and stop scaling when it is not. By computing the γ parameter associated with the different turnover conditions (extracted from data in unlimited environments, **Methods, Figure 3E**), we adapted the model for those different turnover conditions. The model nicely predicted the comet area and bead velocity for the different turnover conditions (**Figure S3D-E**) as well as the progressive decrease of the scaling exponent as a function of the ADF/cofilin concentration (**Figure 3C, red curve**).

**Figure 3.**
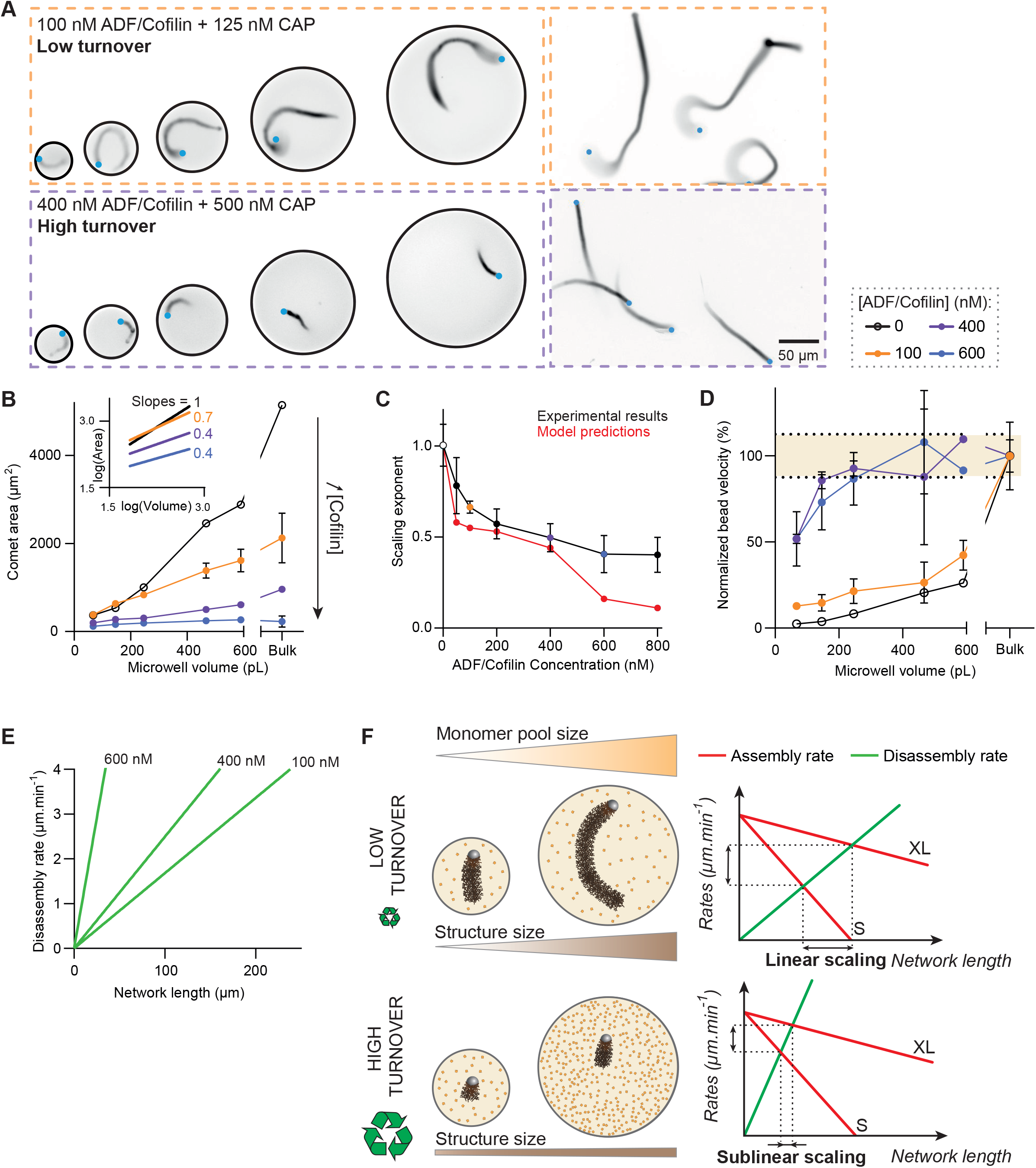
The depletion capacity of growing actin structures determines their scaling abilities. A) Snapshots at 3 hours of actin structures grown in conditions with turnover, in 5 microwell sizes; in grey is actin, in blue the bead. *Biochemical conditions:* 6 µM actin, 12 µM profilin, 60 nM capping protein, 90 nM Arp2/3 complex, 100 nM ADF/cofilin and 125 nM CAP (top row) or 400 nM ADF/cofilin and 500 nM CAP (bottom row). Scale bar: 50 µm. B) Average comet area between 90 and 150 min for each microwell size and bulk at 0, 100, 400 and 600 nM of ADF/cofilin. Inset: corresponding log/log plot showing the scaling exponents. C) Scaling exponents plotted as a function of ADF/cofilin concentration, for all tested conditions (0 to 800 nM ADF/Cofilin). Experimental results (black) are compared against model predictions (red). D) Averaged bead velocity between 90 and 150 min normalized to bulk values, for different conditions (0, 100, 400 and 600 nM ADF/cofilin) for each microwell size. Colored region: ±12.5 % around the reference value (100%). E) Model prediction of the disassembly rate as a function of network length for three different biochemical conditions (100, 400, and 600 nM ADF/Cofilin). F) Scheme and model of the impact of low disassembly and depletion effects on the scaling of structure size: in low turnover, structure size and dynamics scale proportionally to the environment. In high turnover, structure dynamics and size scale sub-linearly with the environment. *Replicates:* see Table

Altogether, these results show that the scalability of a dynamic structure is set by how much it depletes the shared monomer pool (**Figure 3F**), which in turn depends on both the size of the environment and the disassembly rate. When turnover is fast, the disassembly rate matches the assembly rate sooner, so the structure stops growing before it can appreciably deplete the pool; its size becomes largely independent of environment size, and scaling is lost (**Figure 3F, bottom**). When turnover is slow, the pool has time to deplete, so structure size varies with environment size and scaling is preserved (**Figure 3F, top**). In short, fast turnover keeps a structure small enough that it stops depleting the pool, making its size independent of environment size so scaling is lost precisely when turnover is high. In cells, however, multiple structures coexist and draw on the same finite pool. We therefore next introduced several structures into a single environment to ask how limited resources and turnover jointly control the size and dynamics of coexisting structures.

### Increasing the competition between structures is equivalent to tightening the constraints of the environment

To test how competition between structures affects scaling, we turned to the second largest microwells (V = 500 pL, XL microwells) under high-turnover conditions (600 nM ADF/cofilin), where a single actin tail scales significantly sublinearly with the volume, and its size is largely insensitive to the microwell volume. Within these identical wells, we varied the number of beads and asked how each structure’s size changes as more of them compete for the same monomer pool (**Figure 4A, Movie 5**). The effect was clear: comet area decreased as bead number increased, so the more structures shared a well, the shorter each one became (**Figure 4B**). Since the beads share a fixed well volume, adding more of them lowers the volume available to each, so increasing bead number is effectively the same as decreasing the volume per bead. This gives us a natural way to quantify competition: instead of describing each condition by the number of beads, we can describe it by the volume of monomer pool that each structure effectively has to itself. We therefore replotted comet area as a function of volume per bead (**Figure 4C, black curve**). As expected, comet area increases with the volume available per bead, because a structure that competes with fewer neighbors can draw on a larger share of the pool before depleting it.

**Figure 4.**
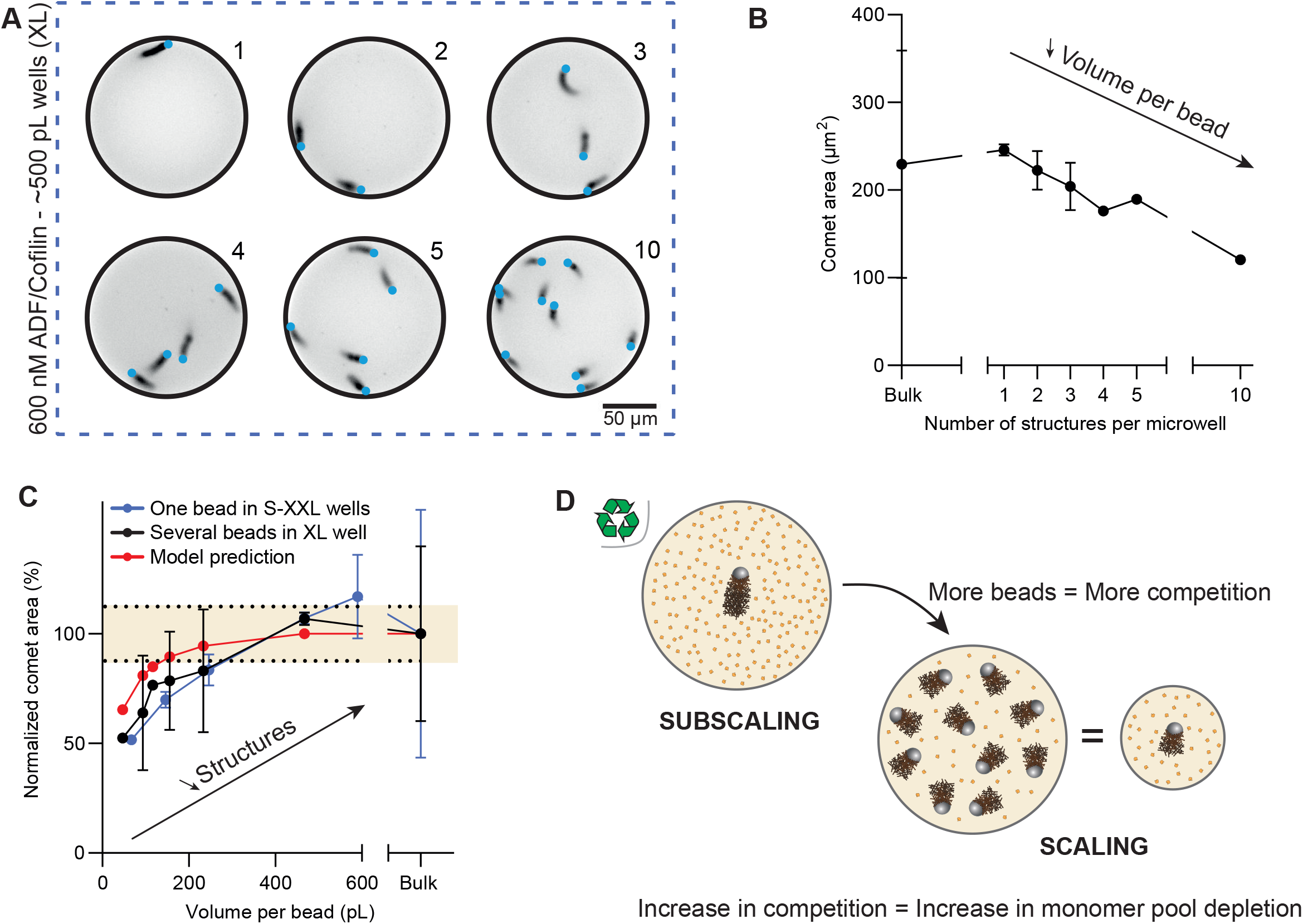
Increased competition among structures is equivalent to stronger environmental constraints. A) Snapshots of actin structures grown in conditions with turnover in microwells of 500 pL (XL) at 3 hours; in grey is actin, in blue the bead. Scale bar: 50 µm. B) Average comet area as a function of number of structures in the microwell. C) Average comet area normalized to bulk value as a function of volume per bead (microwell volume/number of structures) for one bead in several microwell sizes in blue, for several beads in one microwell size in black, and model prediction for several beads in one microwell size in red. D) In sublinear scaling biochemical conditions, the presence of competition in a limited environment is equivalent to an increase in monomer depletion, leading to the scaling of individual structures. *Replicates:* 1 bead: N=3, n=14, 2 beads: N=4, n=14, 3 beads: N=4, n=13, 4 beads: N=3, n=3, 5 beads: N=2, n=16, 10 beas: N=1, n=4, Bulk: N=3, n=413. *Biochemical conditions*: 6 µM actin, 12 µM profilin, 60 nM capping protein, 90 nM Arp2/3 complex, 600 nM ADF/cofilin, 750 nM CAP.

The striking result comes from comparing this curve with a very different experiment: a single bead grown across the full range of microwell volumes (**Figure 4C, blue curve**). The two curves overlap almost perfectly. In other words, several beads in one large well and a single bead in a small well of matching volume-per-bead reach the same size and follow the exact same scaling behavior, under identical biochemical conditions. This behavior is also caught quantitatively by the model (**Figure 4C, red curve**). Competition and confinement are therefore interchangeable: adding a competing structure has the same effect as shrinking the pool available to each one. This equivalence has a direct consequence for scaling. In a fixed environment, where a lone structure would scale sub-linearly, remaining robustly the same size regardless of volume, adding enough competing structures switches the regime back to linear scaling, turning a robust structure into an adaptable one (**Figure 4D**). Competition thus reintroduces the limiting-pool feedback that turnover had removed, allowing structures to once again read out the size of their environment. Overall, these results show that coexisting structures actively contribute to size regulation: by drawing on a shared pool, they constrain one another exactly as a smaller environment would, restoring the scaling that a single dynamic structure had lost.

## Discussion

How subcellular structures acquire a size commensurate with the cell that builds them is a long-standing question in cell biology. Here, by reconstituting the scaling of dynamic cytoskeletal structures from purified proteins, we uncover the physical logic that governs this adaptation. We show that the scalability of an actin network is not a fixed property but an emergent outcome set by three parameters: the turnover rate of the structure, the size of the shared monomer pool, and the size of the environment. Tuning these parameters reveals a continuum of scaling regimes that, together, reconcile the seemingly disparate scaling relationships reported across model systems.

An extreme case is observed without turnover, in which the structure size scales linearly with the environment. In the absence of disassembly machinery, the size of the structure is determined solely by feedback from the limited pool of monomers (Goehring & Hyman, 2012). The limiting pool of components is one of the most used mechanisms to explain intracellular scaling. Larger cells have a higher biosynthetic activity so that they produce more building blocks (Cadart & Heald, 2022; Lin & Amir, 2018; Padovan-Merhar *et al*, 2015). Therefore, intracellular organizations are bigger and scale with cell size. This mechanism has been proposed to explain spindle length (Mitchison *et al*, 2015; Good *et al*, 2013; Hazel *et al*, 2013), microvilli length (Stidwill & Burgess, 1986; Faust *et al*, 2019) or centrosome size regulation (Decker *et al*, 2011).

On the contrary, the introduction of turnover proteins changes the picture entirely. Dynamic structures become highly responsive but lose, to some extent, their capacity to sense the environment: length-dependent disassembly rapidly comes to balance assembly, and in large volumes structure size becomes wholly decoupled from environment size. These observations are reminiscent of what has been reported for the spindle size in large cells of *X. laevis* embryos (Wühr *et al*, 2008) or nucleus in early sea urchin embryos (Mukherjee *et al*, 2020).

The most striking insight lies between these extremes. As turnover increases, the scaling exponent declines smoothly from linear scaling toward sublinear regimes. This single, continuously tunable axis offers a unifying explanation for the wide diversity of scaling laws reported for intracellular organelles across organisms (Marshall, 2020; Chadha *et al*, 2024). Importantly, position along this axis encodes a functional trade-off: a rapidly renewing structure responds rapidly to local perturbation but is largely blind to changes in cell size, whereas a slowly renewing structure is the reverse. In this view, scaling behavior and adaptive strategy are thus two faces of the same physical parameter.

Environmental sensing can nonetheless be regained when multiple structures compete for the same monomer pool. This is physiologically significant, because in many instances it is not the size of a structure but their number that scales with cell size. For example, larger cells assemble more stress fibers and more focal adhesions (Bimbard *et al*, 2025; Alcaraz Casademunt *et al*, 2008; Wada *et al*, 2011; López-Gay *et al*, 2020). Therefore, scaling with increasing the number of structures as a function of cell size is another interesting strategy to adapt to a change in the environment (Marshall, 2020). One of the key aspects of adding multiple structures that coexist in the same environment is that it amplifies the depletion effects on the monomer pool, thereby restoring the feedback mechanism associated with the limited supply of monomers. This resonates with the recent theoretical demonstration that robust size control of multiple structures competing for a finite subunit pool emerges from negative feedback between growth rate and structure size (Banerjee & Banerjee, 2022). Thus, the addition of multiple competing structures that will deplete the monomer pool is a way to ensure that this feedback is present and robustly determines the size of the structures.

Extending these principles to actin structures in living cells raises clear challenges, each of which defines a concrete future direction. First, the cytoplasmic monomer pool is not homogeneous, and functionally distinct pools of actin monomers can coexist within the cell (Skruber et al, 2018; Galbraith et al, 2026; Vedula et al, 2021; Lee et al, 2013; Vitriol et al, 2015; Fan et al, 2012). In its current form, our biomimetic system has a common but homogeneous pool of resources. Introducing controlled heterogeneity and letting structures with different architecture and turnover compete within one environment will test how far these principles hold as complexity increases. Second, quantifying the size of individual actin structures in cells is difficult because they are physically interconnected. New image analysis tools (notably based on AI, Aljapur et al, 2026; Edwards et al, 2021; Rostami et al, 2023) now makes it realistic to measure architecture-specific size as a function of cell size. In addition, correlating this size to the available pool of actin monomers also poses a challenge when it comes to measure the quantity of G-actin in cells (both at the whole-cell scale and at the local scale, in the cytoplasm, (Skruber et al, 2018). The development of new tools (FRET sensors e.g. (Fedoryshchak et al, 2026)) will be of a primary importance in order to correlate the size of actin structures with the local availability of actin monomers.

Overall, these findings establish a mechanistic framework for the scaling of dynamic cytoskeletal structures. This framework establishes that their ability to adapt to their environment is set by the degree to which a structure depletes shared resources as it grows. Although derived from a biomimetic actin system, the principle is generic: it should apply not only to other polymers but to any dynamic cellular organelle whose assembly and disassembly rates respond to environmental cues (Mohapatra *et al*, 2016). In reducing the diversity of biological scaling laws to a small set of physical parameters, our work offers a quantitative foundation for understanding how cells match the size of their internal architecture to their own.

## Acknowledgments

We thank all the members of Cytomorpholab for insightful discussions. This work was supported by the grant ANR-24-CE13-3582 SCALING to A.Co. This work also received funding (PhD grant for A. Ca and funding for MuLife Imaging platform) from GRAL, the Grenoble Alliance for Integrated Structural and Cell Biology, a program of the Chemistry Biology Health Graduate School of Université Grenoble Alpes (ANR-17-EURE-0003) and from the CNRS through the MITI interdisciplinary programs.

## Author contributions

Conceptualization: A.Co, A.Ca, LB, MT / Experiments: A.Ca, CG, NM / Methodology: A.Ca, BV / Data Analysis: A.Ca / Mathematical modeling: AM / Funding acquisition: A.Co / Writing of the paper: A.Co, A.Ca

## Figures legends

**Figure S1.**
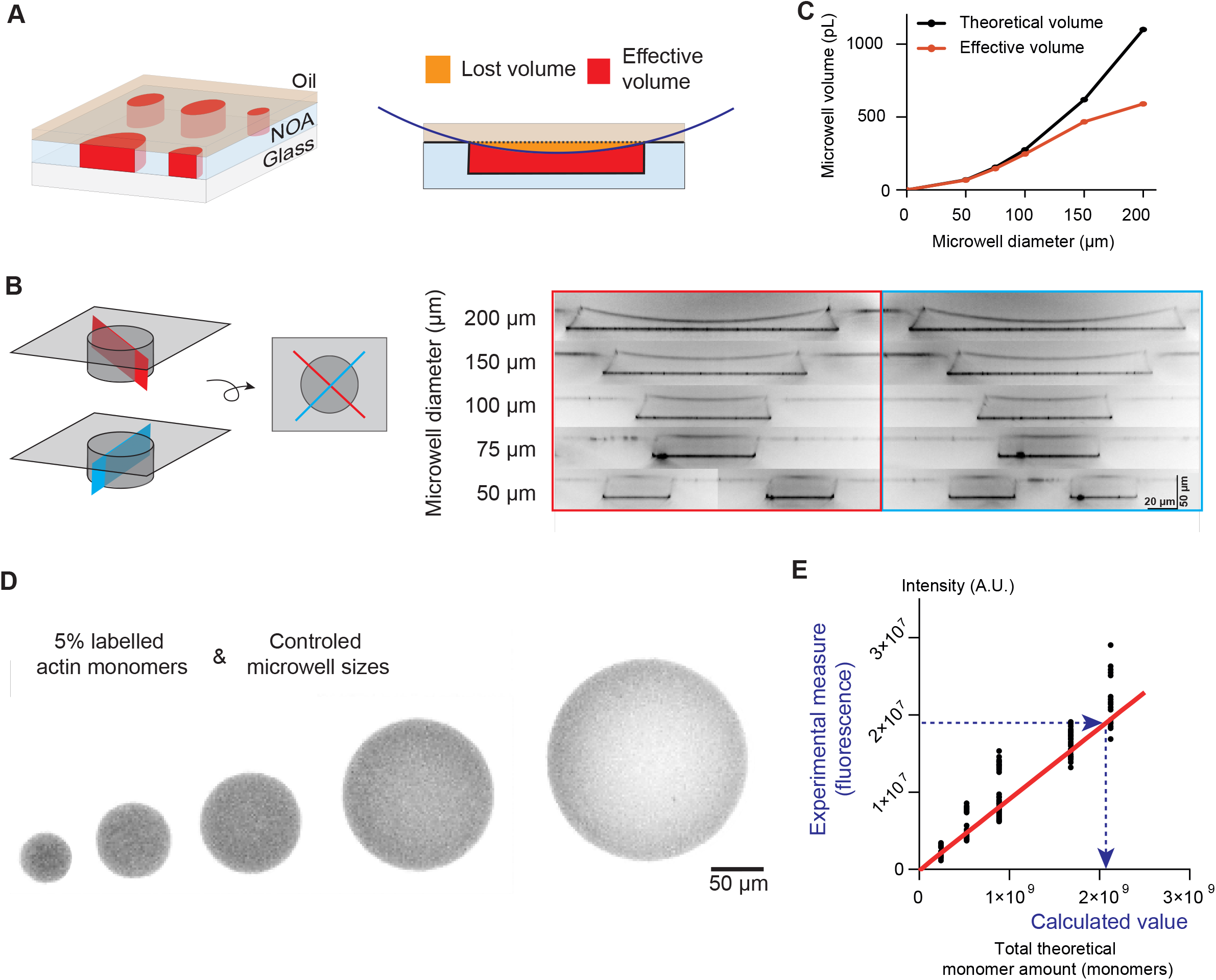
Measure and calibration of all microwell sizes. A)Scheme of the microwell environment: detail of the volume lost to the oil meniscus on the top layer. Spinning disk Z-stacks of the lipid layer in each microwell size, 2 perpendicular stacks (plane shown on scheme, in blue and red) per microwell were performed, showing the oil meniscus at the top of each microwell. Scale bar: 50 µm vertical, 20 µm horizontal. C) Plot of the theoretical and effective volume for each microwell size, after lost volume calculation. D) Snapshot of actin filled microwells, from which intensity measurements were extracted. Scale bar: 50 µm. E) Calibration curve plot of measured intensity as a function of theoretical actin monomer quantity in each microwell size. Slope= 0.010, R^2^= 0.92.

**Figure S2.**
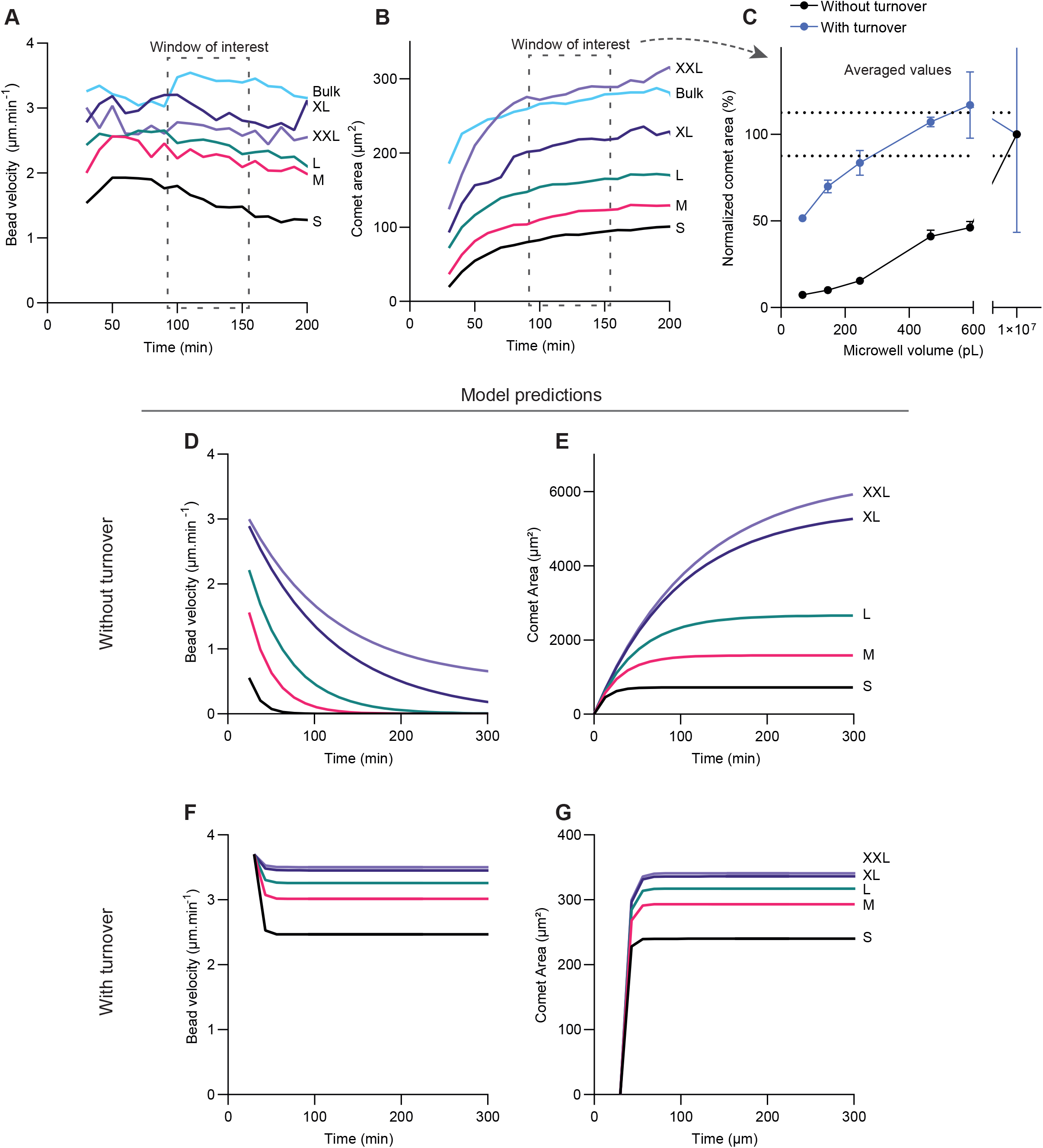
Overtime measures of actin comets with turnover at 600 nM ADF/Cofilin and 750 nM CAP and model predictions. A) Bead velocity overtime for each microwell size and bulk, corresponding to the snapshots from **Figure 2A**. Window of interest highlight which data was used in the normalized plot in **Figure 2B**. B) Comet area overtime for each microwell size and bulk, corresponding to the snapshots from **Figure 2A**. Window of interest highlight which data was used in the normalized plot in panel C. C) Normalized averaged comet area for each microwell size and bulk. D) Model predictions of bead velocity overtime in conditions without turnover for each microwell size (relative to **Figure 1G**). E) Model prediction of comet area overtime in conditions without turnover for each microwell size (relative to **Figure 1E**). F) Model predictions of bead velocity overtime in conditions with turnover for each microwell size (relative to **Figure S2A**). G) Model prediction of comet area overtime in conditions with turnover for each microwell size (relative to **Figure S2B**).

**Figure S3.**
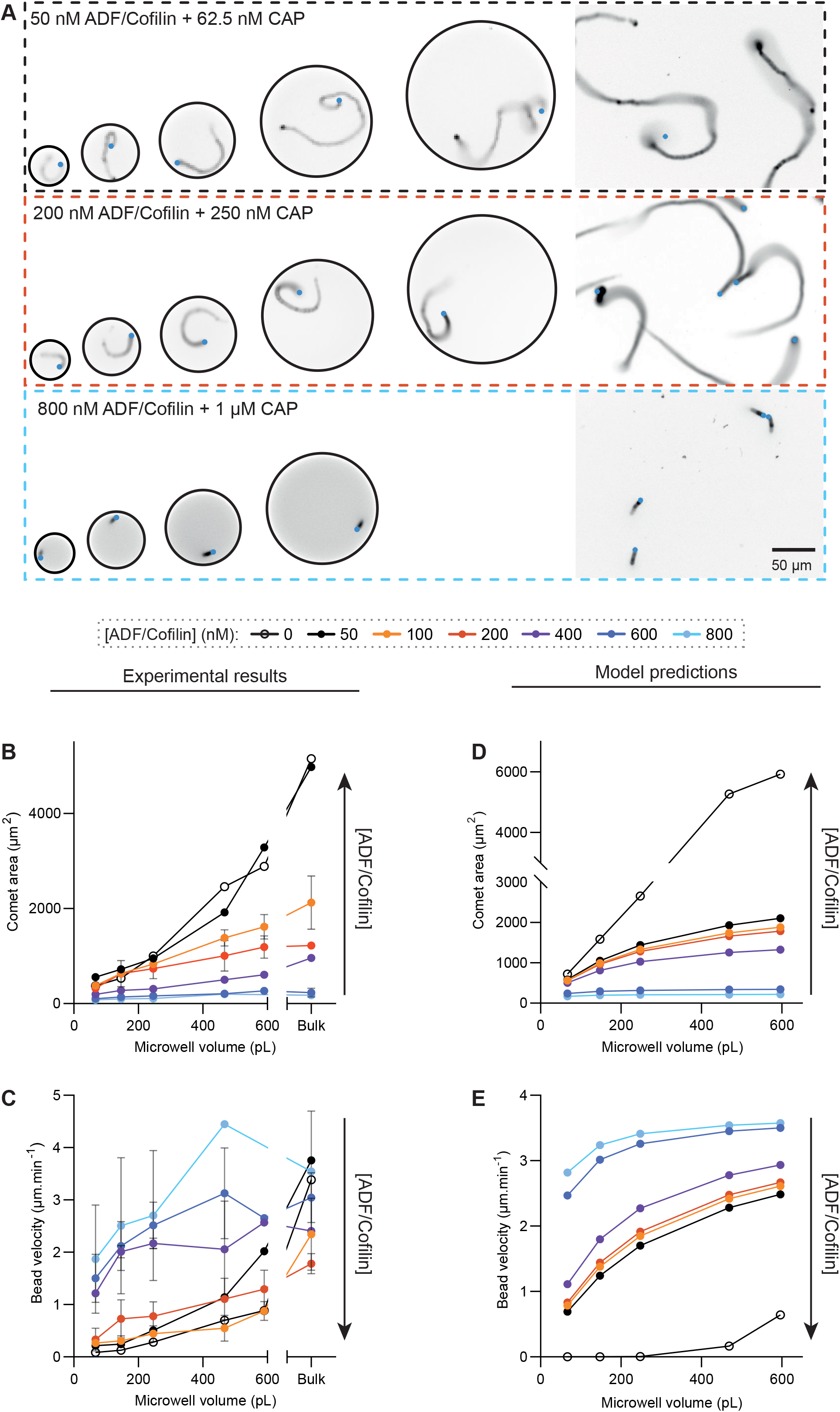
Additional results and model predictions for different turnover conditions. A) Snapshots at 3 hours of actin structures in conditions with turnover in 5 microwell sizes and in bulk; in grey is actin, in blue the bead. *Biochemical conditions:* 6 µM actin, 12 µM profilin, 60 nM capping protein, 90 nM Arp2/3 complex, 50, 200 and 800 nM ADF/cofilin and 62.5 nM CAP, 250 and 1000 nM CAP respectively. Scale bar: 50 µm. B) Average comet area between 90 and 150 min for each microwell size and bulk, in all biochemical conditions: 0, 50, 100, 200, 400, 600, and 800 nM ADF/Cofilin and corresponding CAP. C) Average bead velocity between 90 and 150 min for each microwell size and bulk, in all biochemical conditions. D) Model predictions for comet area across microwell volumes for all biochemical conditions. E) Model predictions for bead velocity across microwell volumes for all biochemical conditions. *Replicates:* See Table.

## Replicates Table

| ADF/cofilin concentration | 50 nM |  | 100 nM |  | 200 nM |  | 400 nM |  | 600 nM |  | 800 nM |  |
|---|---|---|---|---|---|---|---|---|---|---|---|---|
|  | N | n | N | n | N | n | N | n | N | n | N | n |
| S | 1 | 2 | 3 | 17 | 4 | 55 | 2 | 47 | 4 | 76 | 2 | 43 |
| M | 1 | 6 | 4 | 25 | 3 | 43 | 2 | 29 | 4 | 80 | 2 | 37 |
| L | 1 | 13 | 4 | 59 | 4 | 59 | 2 | 24 | 4 | 83 | 2 | 36 |
| XL | 1 | 6 | 4 | 22 | 4 | 18 | 2 | 10 | 3 | 14 | 0 | 0 |
| XXL | 1 | 1 | 4 | 19 | 3 | 11 | 1 | 5 | 3 | 12 | 0 | 0 |
| Bulk | 1 | 164 | 4 | 582 | 2 | 156 | 1 | 146 | 3 | 413 | 2 | 438 |

## Movies legends

**Movie 1: Actin comet assembly without turnover in microwells of 5 different sizes and in bulk, related to Figure 1D**. Time lapse of 5 microwells of different sizes and bulk with one actin comet each, in conditions without turnover. Beads are shown in blue, labeled actin in grey. Images were acquired every 10 min over 4:30 hours. The movie was compressed in JPEG at 8 frames per second. Scale bar: 50 µm.

**Movie 2: Actin comet assembly with turnover at 600 nM ADF/Cofilin and 750 nM CAP, in microwells of 5 different sizes and in bulk, related to Figure 2A**. Time lapse of 5 microwells of different sizes with one actin comet each, and representative portion of a field of view in bulk, in conditions with turnover. Beads are shown in blue, labeled actin in grey. Images were acquired every 10 min over 2:30 hours. The movie was compressed in JPEG at 8 frames per second. Scale bar: 50 µm.

**Movie 3: Actin comet assembly with turnover at 100 nM ADF/Cofilin and 125 nM CAP, in microwells of 5 different sizes and in bulk, related to Figure 3A**. Time lapse of 5 microwells of different sizes with one actin comet each, and representative portion of a field of view in bulk, in conditions with turnover. Beads are shown in blue, labeled actin in grey. Images were acquired every 10 min over 5:10 hours. The movie was compressed in JPEG at 8 frames per second. Scale bar: 50 µm.

**Movie 4: Actin comet assembly with turnover at 400 nM ADF/Cofilin and 500 nM CAP, in microwells of 5 different sizes and in bulk, related to Figure 3A**. Time lapse of 5 microwells of different sizes with one actin comet each, and representative portion of a field of view in bulk, in conditions with turnover. Beads are shown in blue, labeled actin in grey. Images were acquired every 10 min over 4:10 hours. The movie was compressed in JPEG at 8 frames per second. Scale bar: 50 µm.

**Movie 5: Different numbers of actin comet assembly with turnover at 600 nM ADF/Cofilin and 750 nM CAP, in microwells of 500 pL (XL), related to Figure 4A**. Time lapse of microwells with increasing numbers of beads, in conditions with turnover. Beads are shown in blue, labeled actin in grey. Images were acquired every 10 min over 2:40 hours. The movie was compressed in JPEG at 8 frames per second. Scale bar: 50 µm.

## Material and methods

### Mathematical Model

#### Quantitative model

We use the following model to describe the comet sizes and velocities. Let *L* be comet length. The rate of the length change, *dL/dT*, is the difference of the growth comet rate (which is the same as the bead velocity, *V*) and of the comet disassembly rate: *dL*/*dT* = *V* − *D*, where *D* is the disassembly rate. Previous studies established that *V* = *kG*, where *G* is the G-actin concentration in the well (Boujemaa-Paterski *et al*, 2017; Colin *et al*, 2023; Guérin *et al*, 2025).

Here *k* = *k*_*on*_*δ*Φ where *k*_*on*_*δ* ≈ 30 nm.sec^-1^.*µ*M^-1^ (Boujemaa-Paterski *et al*, 2017) and factor Φ ≈ 0.3 is responsible for mechanical and geometric effects of skewed and loaded actin filament elongation kinetics at the bead surface (Colin *et al*, 2023).

In (Manhart *et al*, 2019), we showed that the comet disassembly takes place at the distal end of the comet according to the following kinetics. After the actin network is assembled at the surface of the bead, it takes time τ until the network ‘ages’ to the point of crumbling apart (this aging is due to loosening connectivity in the network). Time τ is the function of cofilin concentration (Manhart *et al*, 2019). In the case of the steady motility of the bead with velocity *V, d* = *Vτ*. Let us compare this simple result with the general kinetic equation *dL*/*dT* = *V* − *D*. According to this equation, in the steady state, *D* = *V*. If the net disassembly rate *D* is proportional to the comet length *L*: *D* = *γL*, where *γ* is the proportionality coefficient, then *γL* = *V*, and comparing this expression with *L* = *Vτ*, we see that *γ* = 1/*τ*. Thus, we use the simplest possible model: *dL*/*dT* = *V* – *γL* where parameter *γ* is a constant dependent on the cofilin concentration. In (Manhart *et al*, 2019), we demonstrated both experimentally and mathematically that for longer comets, this simplest model is indeed the case, while for shorter comets, the net disassembly rate is proportional to ~*L*^2^ due to the effect of the gradual accumulation of cofilin in the comet. Other theoretical model also proposed the dependence

~*L*^2^ of the net disassembly rate (Mohapatra *et al*, 2016). We simulated the model in both of these cases, found that results differ relatively little, and chose the simplest model.

To find the G-actin concentration in the microwell, we use the following simple mass balance argument. First, for simplicity, we do not separately account for ATP- and ADP-G-actin. Second, introducing a constant parameter *β* that converts a unit length of the comet into the number of monomers, we have 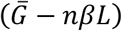 monomers in the microwell, where *n* is the number of the beads and 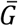 is the total number of monomers in the microwell initially, before the polymerization started. Note that if *W* is the volume of the well, then 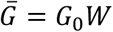, where *k*_0_ = 6 *µM* is the initial actin concentration.

Thus, 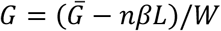. Collecting two principal equations, the model is:

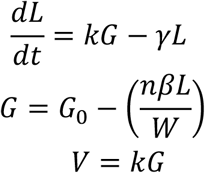

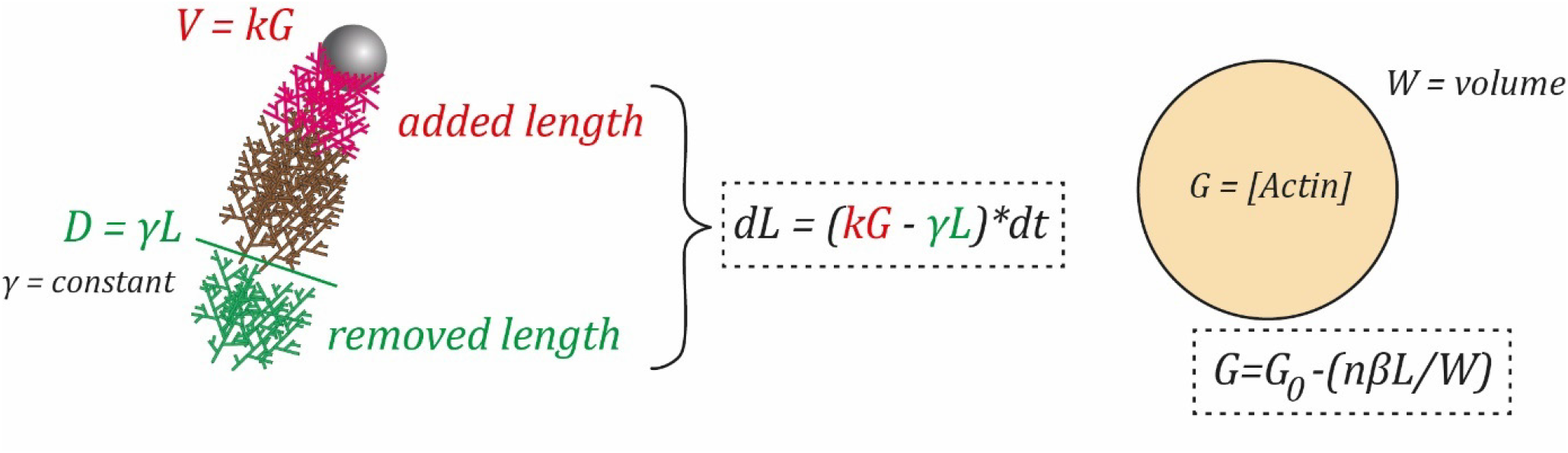

#### Model parameters

We find the model parameters as follows:

Parameter *k*: estimated above, *k* = 0.01 *µm. s*^−1^. *µM*^−1^

Parameter *γ*: in the presence of cofilin, when beads are moving steadily:

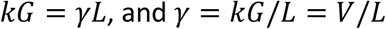

We have measured average bead velocities and comet lengths for all used cofilin concentrations *in the unlimited pool* and found respective disassembly rates for all these conditions. Specifically, *γ* = (0.013, 0.015, 0.016, 0.024, 0.11, 0.18) min^-1^ for cofilin concentrations C = (50, 100, 200, 400, 600, 800) nM, respectively.

Parameter G_0_: *k*_0_ = 6 *µM*

Parameter *<u>W</u>*: we used five measured volumes for S, M, L, XL and XXL wells (**Figure S1**).

Parameter *β*: we used the final comet length, 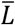, measured in the experiment without turnover in the largest microwell with volume *W*_*XXL*_, as well as the measured fraction of the amount of actin remaining unpolymerized in this condition, *f*_*XL*_. Then:

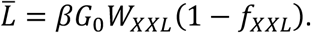

We calculated 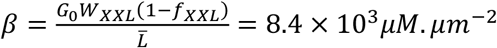

Finally, when we convert the predicted comet length into the comet area, we multiplied the length by the width; the latter was taken to be ~ 15 *µm* based on analysis of the images.

Note that experimental data from smaller wells under the same conditions give close estimates. We chose the data from the largest well because of potentially smallest random effects from larger numbers of molecules involved in the actin dynamics in the largest limited pool.

We emphasize therefore that we calibrated the model using only the data from (i) no cofilin in the unlimited pool and one limited pool, (ii) various cofilin concentrations in the unlimited pool, in the case of a single bead. Then, we predict the comet sizes and bead velocities for various cofilin concentrations in the limited pools and for various number of beads without further fitting (without free parameters).

#### Scaling and non-dimensionalization

Characteristic actin concentration is *G*_0_.

Characteristic convenient length scale in this model is 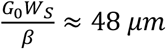, where *W*_*s*_ is the volume of the smallest well. Characteristic convenient time scale in this model is 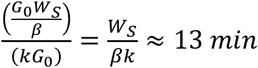. Using these scales, we introduce the non-dimensional actin concentration, *g*, comet length, *l*, velocity, *v*, and time *t*. The non-dimensionalized equations have the form:

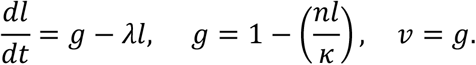

Thus, the predicted actin dynamics in our system depends on just two non-dimensional parameter combinations:

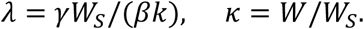

Note that parameter *λ* is the cofilin-dependent disassembly rate multiplied by the characteristic time scale, while parameter *k* is the ratio of the volume of the well and volume of the smallest well.

Non-dimensional model equations can be solved analytically (see below). We convert the dimensionless solutions for the comet length by multiplying by the length scale, and for the bead velocity by multiplying by the length scale and dividing by the time scale.

#### Model predictions for the assembly case

When cofilin is absent, there is no comet disassembly, and the model becomes:

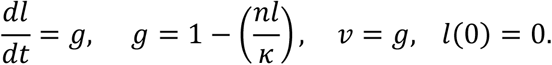

For a single bead, the solutions of these equations are:

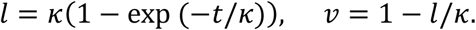

#### Model predictions for the turnover case

In the case of steady beads’ motility,

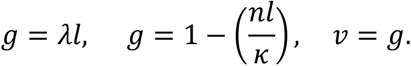

The solutions of these equations are:

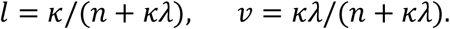

Full time dependent solutions for any bead number are:

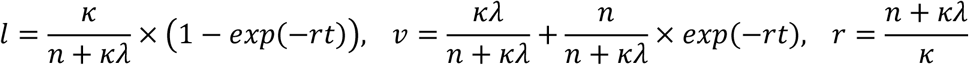

### Protein purification

#### Actin

Fibric actin was rehydrated from rabbit skeletal-muscle acetonic powder 2 times in G-Buffer (2 mM Tris pH=8, 0.5 mM ATP, 0.2 mM DTT, 0.1 mM CaCl_2_, 1 mM NaNH_3_) before centrifugation (16k rpm, 30 min, 4°C, JA25.50 Beckman rotor), and supernatant gathered and filtered over glass wool. Actin-G was polymerized by adding 50 mM KCl and 2 mM MgCl_2_ under agitation for 1 hour, then 0.8 M KCl is added as powder for tropomyosin dissociation from F-actin. The solution was centrifuged (35k rpm, 2 hours, 4°C, TI45 Beckman rotor), and the fibers of the resuspended pellet in G-Buffer were broken through a dounce homogeneizer. The resulting actin gel was dialyzed (12-14 kDa) in G-Buffer over two days at 4°C for depolymerization. Dialysate was then centrifuged (2 hours, 75k rpm, 4°C) to gather monomeric actin, which was loaded on Sephacryl S300-26-600 gel filtration column before being stored in 2 mL fractions at 4°C, used for up to 3 weeks.

A portion of the purified actin was labelled as follows. Purified G-actin was dialyzed in marking buffer (20 mM Hepes pH=7.8, 50 mM KCl, 0.1 mM CaCl_2_, 0.2 mM ATP) and left overnight. 0.5 mg of Alexa568 in 30 µL DMF was added per 2 mL of actin and stirred for 1 hour at RT in the dark. Solution was centrifuged (85k rpm 45 min, 4°C) and the pellet resuspended in G-buffer, before dialysis in G-buffer (3 hours, 4°C). The labelled actin was loaded on G25 column and the first actin fractions were pooled together. 0.1 M KCl and 2 mM MgCl_2_ were added and the solution left for polymerization to stir for 1 hour at RT in the dark. Solution was again centrifuged (85k rpm, 45 min, 4°C, TLA110 Beckman rotor), resuspended, and dialyzed overnight to trigger depolymerization. Finally, labelled monomeric actin was centrifuged and stored in 60% glycerol at −20°C and used for up to 3 weeks.

#### Profilin

Human profilin (HP) was expressed in *Escherichia coli* strain Rosetta 2 (DE3) pLysS. Cells were cultured in 2 L LB at 37°C for 4h. Pelleted cells where resuspended in 20 mM Tris pH 8, 20 mM KCl, 1 mM EDTA, 1 mM DTT, 2 M Urea, and lysed by sonication. The supernatant, clarified by centrifugation (16 000 rpm, 30 min, 4°C), was loaded on a poly-L-proline Sepharose column and washed with the same buffer. Protein elution was followed using Bradford method, then completed with 8 M Urea. The fractions of interest were pooled and dialyzed (3.5 kDa) in 2 L storage buffer (20 mM Tris pH8, 1 mM EDTA, 1 mM DTT) at 4°C. Dialyses were repeated for a week with a change of bath 2 times a day. Protein was centrifuged (55k rpm, 30 min, 4°C) before being flash frozen and stored at −80°C.

#### WA (from NWASP)

Snap-Streptavidin-WA-His (pETplasmid) was expressed in Rosetta 2 (DE3) pLysS (Merck, 71403). Culture was grown in TB, induced with 0.5 mM IPTG at 16°C overnight. Cells were pelleted and resuspended in lysis buffer (20 mM Tris pH8, 500 mM NaCl, 1 mM EDTA, 15 mM Imidazole, 0,1% TritonX100, 5% Glycerol, 1 mM DTT, proteases inhibitors) before sonication and centrifugation (16K rpm, 30 min, 4°C). Supernatant was loaded on Ni Sepharose high performance resin (GE Healthcare Life Sciences, ref 17526802) and kept under agitation for 2 hours at 4°C. Resin was loaded on a column and washed repeatedly (20 mM Tris pH8, 500 mM NaCl, 1 mM EDTA, 30 mM Imidazole, 1 mM DTT) before elution with 300 mM Imidazole. Eluted protein was then dialyzed overnight (20 mM Tris pH8, 150 mM NaCl, 1 mM EDTA, 1 mM DTT) before centrifugation (65k rpm, 30 min) and load on S200-16-600 gel filtration column. Finally, protein was concentrated with Amicon Ultra-15 10K, centrifuged at 3500g, 10 min, before flash freezing and storage at −80°C.

#### Arp2/3 complex

The Arp2/3 complex was purified from bovine thymus. Calf thymus was cut into approximately 1 cm pieces and mixed with extraction buffer (20 mM Tris pH 7.5, 25 mM KCl, 1 mM MgCl_2_, 5% glycerol, protease inhibitors) for 1-2 minutes, then shaken in a beaker for 30 minutes. The extract was then centrifuged in a benchtop centrifuge at 4.5k RPM for 5 minutes, and the supernatant clarified at 18k rpm for 25 minutes at 4°C. The supernatant was then filtered over glass wool, the pH was fixed at 7.5 with KOH and centrifugation was carried out at 38k RPM at 4°C for 1 hour. The medium aqueous phase was transferred to a chilled glass beaker, the extract was precipitated with 50% ammonium sulfate, let for stirring at 4°C for additional 30min, and centrifuged at 18k RPM for 30 minutes at 4°C. The pellet was resuspended in 10 mL (for 200g thymus) extraction buffer + 0.2 mM ATP, 1 mM DTT and protease inhibitor. It was then dialyzed overnight in Arp2/3 dialysis buffer (20 mM Tris pH 7.5, 25 mM KCl, 1 mM MgCl_2_, 5% glycerol, 1 mM DTT and 0.2 mM ATP). A GST-WA glutathione sepharose column was prepared and washed with extraction buffer containing 0.2 mM ATP, 1 mM DTT and protease inhibitors. The dialyzed extract was passed over the GST-WA column. Next, the column was washed with 20 mL extraction buffer with 0.2 mM ATP, 1 mM DTT and then with 20 mL extraction buffer with 0.2 mM ATP, 1 mM DTT and 100 mM KCl. The Arp2/3 complex was eluted with 20 mL extraction buffer with 0.2 mM ATP, 1 mM DTT and 200 mM MgCl_2_, then dialyzed in source buffer A (piperazine-N,N0-bis(2-ethanesulfonic acid) (PIPES) pH 6. 8, 25 mM KCl, 0.2 mM ethylene glycol-bis(b-aminoethyl ether)-N,N,N0,N0-tetraacetic acid (EGTA), 0.2 mM MgCl_2_ and 1 mM DTT) overnight. The Arp2/3 complex was then loaded onto a MonoS column and eluted with source buffer B (piperazine-N,N0-bis(2-ethanesulfonic acid) (PIPES) pH 6.8, 1 M KCl, 0.2 mM ethylene glycol-bis(b-aminoethyl ether)-N,N,N0,N0-tetraacetic acid (EGTA), 0.2 mM MgCl_2_ and 1 mM DTT). The Arp2/3 complex was dialyzed in storage buffer (10 mM Imidazole pH 7.0, 50 mM KCl, 1 mM MgCl_2_, 0.2 mM ATP, 1 mM DTT and 5% glycerol), flash-frozen in liquid nitrogen and stored at −80°C.

#### Capping Protein

Mouse Capping Protein (CP) was expressed in *Escherichia coli* strain Rosetta 2 (DE3) pLysS. Cells were cultured in 2 L LB at 24°C overnight. Cells were pelleted and resuspended in lysis buffer (20 mM Tris pH 8.0, 250 mM NaCl, 10 mM Imidazole, 5% Glycerol, 1 mM DTT, 1 mM EDTA) + EDTA-free protease inhibitors cocktail tablet. Cells were sonicated and centrifuged at 16 000 rpm for 30 min, 4°C. Supernatant was applied to 1 mL of Ni sepharose fast flow resin (GE healthcare). After one hour at 4°C under gentle rotation, resin was washed with 20 volumes of Buffer A (20 mM Tris pH8, 250 mM NaCl, 1 mM EDTA, 1 mM DTT, 20 mM Imidazole). Elution was performed with buffer A + 300 mM Imidazole. Purified protein was dialyzed over night against a storage buffer (20 mM Tris pH 8.0, 1 mM DTT, 1 mM EDTA, 0.2 mM CaCl_2_), then centrifuged (50k rpm, 25 min, 4°C) to remove aggregates. Aliquots were flashfrozen in liquid nitrogen and stored at −80°C.

#### ADF/Cofilin

D34C mutant *S. cerevisiae* ADF/cofilin was expressed in *Escherichia coli* strain Rosetta 2 (DE3) pLysS then purified as follows. Cells were pelleted then resuspended in 40 mL lysis buffer (25 mM Tris pH 8, 300 mM NaCl, 1 mM EDTA, 1 mM TCEP, 0.1% Triton, 5% Gycerol, EDTA-free proteases inhibitors). The cell solution was sonicated for 2×2min before centrifugation (16k rpm, 30min, 4°C). Supernatant was loaded on a Glutathione beads resin and kept under agitation for 1h30 at 4°C before centrifugation (100g, 1min, 4°C). The resin was loaded on a column and washed with TBSE buffer (25 mM Tris pH 8, 300 mM NaCl, 1 mM EDTA, 1 mM TCEP). GST site was cleaved with Thrombin in 4 mL TBSE buffer under agitation for 2 hours at 4°C, and protein was eluted on a column with TBSE buffer. Finally, Thrombin was removed with para aminobenzamidine with agitation for 10 min at 4°C before cofilin final elution and dialysis overnight (MWCO 3500 kDa). Part of the resulting protein content was labelled by adding 5 excesses of alexa-488 C5-maleimide (60 µL at 10 mM) + 1 mL of cofilin in TCEP at 100 µM overnight with agitation at 4°C, and following elution on Sephadex G25 Gel Filtration. After reconcentration with Amicon Ultra-15 3.5 kDa cut-off, aggregates were removed with centrifugation (55k rpm, 30 min, 4°C). Aliquots were flash frozen in liquid nitrogen and stored at −80°C.

#### CAP

The CAP1 protein was expressed in Arctic *Escherichia coli* (Sigma) in LB medium at 13°C for 30 hours. The bacteria were pelleted and resuspended in buffer A (50 mM Tris pH 7.5, 500 mM NaCl, 25 mM imidazole) and lysed by sonification in the presence of protease inhibitors (EDTA-free tablets: 200 mg/ml PMSF, 1 mg/ml leupeptin, 1 mg/ml aprotinin, 1 mg/ml pepstatin A, 150 mg/ml benzamidine hydrochloride, all from Sigma-Aldrich). The supernatant, clarified by centrifugation (16 000 rpm, 30 min, 4°C), was then loaded to a 5 mL HisTrap Ni-NTA column coupled to AKTA Pure protein purification system (GE Healthcare). The His-SUMO-tagged CAP1 protein was eluted from the nickel column with a gradient up to 250 mM imidazole. The His-SUMO-tag was then cleaved from the CAP protein in the presence of SENP2 protease (His-Tagged). The cleaved CAP was separated from the uncleaved protein and the protease by collecting the flowthrough after loading on another HisTrap Ni-NTA column, after which the cleaved protein was subjected to gel filtration runs by using Superdex S200-26-600 gel filtration column equilibrated in 20 mM Tris pH 7.5, 150 mM NaCl, 1 mM DTT, 1 mM EDTA. Peak fractions from the same elution volume were combined, and concentrated with Amicon Ultra-15 30 kDa cut-off centrifugal filter device. Aggregates were removed with centrifugation (30k rpm, 25 min, 4°C), and aliquots were flash frozen in liquid nitrogen and stored at −80°C.

### Microwells preparation

Microwells used in this study have diameters of 50, 75, 100, 150, and 200 µm and a height of 35 µm, giving them an effective volume of about 67, 146, 246, 467, and 590 pL respectively (**Figure S1**). All steps of microfabrication followed classical protocols described in (Vianay *et al*, 2025). SU8 mold with pillars was prepared using standard protocols and silanized with Trichloro(1H,1H,2H,2H-perfluoro-octyl)silane for 1 hour and heated for 1 hour at 120°C. From the SU8 mold, a PDMS primary mold was prepared (Dow, SYLGARD 184 silicone elastomer kit) with a 1:10 w/w ratio of curing agent. PDMS was cured at 70°C for at least 2 hours. PDMS primary mold was then silanized with Trichloro(1H,1H,2H,2H-perfluoro-octyl)silane for 1 hour and heated for 2 hours at 100°C. PDMS was then poured on top of the PDMS primary mold to prepare the PDMS stamps. The first set of PDMS stamps were used to prepare an Epoxy mold prepared with a ratio of 6:10 in weight (Epoxy resin R123, Soloplast).

Coverslips were cleaned with the following protocol: they were first wiped with ethanol (96%) then sonicated 30 minutes in Hellmanex 2% at 60°C. After the sonication, coverslips were rinsed in several volumes of mqH_2_O and kept in water until use. Just before use, coverslips were dried with compressed air. For the microwells preparation, PDMS Stamps were cut in pieces and placed on the coverslips with the pillars facing the coverslip. A drop (5 µL) of NOA 81 (Norland Products) was then placed on the side of the PDMS stamp and NOA was allowed to go through the PDMS Stamp by capillarity. When the NOA filled all the stamp, it was polymerized with UV light for 12 minutes (UV KUB2/ KLOE; 100% power). After polymerization of the NOA, PDMS stamp was removed and the excess of NOA was cut. Then, an additional UV exposure of 2 minutes was done and the microwells were placed on a hot plate at 110°C for at least 3 hours (or at 60°C overnight) to tightly bind the NOA to the glass. Steps from coverslips cleaning to the end are done in a single day.

### Measure of microwells effective volume

As we are using a large range of microwell sizes, the oil layer used to close the wells can affect the effective volumes of our microwells (through meniscus effects, described in (Vianay *et al*, 2025)). In order to precisely estimate microwells volume, we made Z-stack of each size of microwells using a confocal spinning-disc system (Eclipse TiE Nikon inverted microscope equipped with a CSUX1-A1 Yokogawa confocal head, an Evolve EMCCD camera (Photometrics), and a Plan Fluor 40X objective). Z-stacks were acquired with Metamorph software (Universal Imaging). FiJi (Schindelin *et al*, 2012) software was then used to create stacks of each microwell size and measure curvature of the oil layer at the top of the wells (as described in (Vianay *et al*, 2025)). We verified that this curvature was identical across all well sizes. This allowed us to then calculate the effective volume for each microwell size (**Figure S1**), which were then used in all data processes as the standard volume measurements.

### Lipids/SUV preparation

For regular microwell passivation, L-a-phosphatidylcholine (EggPC, Avanti, 840051C, 10 mg/mL) was used. The SUVs were prepared as follows: 100 µL of lipids were introduced in a glass tube and the mixture was dried with nitrogen gas. The dried lipids were incubated in a vacuum overnight. After that, the lipids were hydrated in the SUV buffer (10 mM Tris (pH 7.4), 150 mM NaCl, 2 mM CaCl_2_). The mixture was sonicated on ice for 10 minutes. The mixture was then centrifuged for 10 min at 20,238 xg to remove large structures. The supernatants were collected, stored at 4°C and used for up to 3 weeks.

### SilanePEG30k slides

SilanePEG (mPEG-Silane, MW 30 kDa, PSB-2014, Creative PEG works) solution was prepared at a final concentration of 1 mg/mL in 96% ethanol and 0.1%(v/v) HCl. Slides were cleaned with the following protocol: they were wiped with ethanol (96%) then sonicated for 30 minutes at 60°C in Hellmanex 2%. They were then rinsed with several volumes of mqH2O and dried with compressed air. Slides were then plasma cleaned for 5 minutes at 80% power and directly immersed in the silanePEG solution. They were kept in the silanePEG solution until use.

### Polystyrene beads coating

Polystyrene beads coating with NPF was done following classical protocols (Reymann *et al*, 2011). 20 µL of 4.5 µm polystyrene beads (Fluoresbrite BB Carboxylate Microspheres, 2.5% solids, 4.5 µm in diameter) were centrifuged at 13, 000 xg for 2 minutes on a mini spin plus Eppendorf centrifuge (Rotor F45-12-11). The pellet was then resuspended in 200 µL of a 400 nM NPF (Snap-Streptavidin-WA-His) solution. Beads were incubated for 15 minutes at 15°C at 950 rpm in a thermoshaker. They were then centrifuged for 2 minutes at 3800 xg, resuspended in 200 µL of BSA 1% and let on ice for 5 minutes. Beads were finally centrifuged again 2 minutes at 3800 xg and resuspended in 50 µL of BSA 0.1%. The bead coating was redone every day.

### Bead motility assay in microwells

A typical experiment of bead motility in microwells was performed as follows. The coverslip with microwells was activated with plasma for 2 minutes at 80% power. Just after the plasma, either a PDMS chamber or a slide passivated with SilanePEG 30k and 180 µm height double-side tape was mounted with the microwells coverslip. Lipids were rapidly inserted in the chamber and incubated for 10 minutes. Lipids were then rinsed with 1 mL of SUV buffer (5×200 µL) and 200 µL of 1x HKEM buffer (50 mM KCl, 15 mM HEPES pH=7.5, 5 mM MgCl_2_, 1 mM EGTA). The reaction mix with the different proteins was then prepared and flowed in the flow cell. A typical reaction mix (60 µL) was prepared with beads coated with WA (activator of the Arp2/3 complex) and 6 µM of actin monomers, 12 µM profilin, 90 nM Arp2/3 complex, 40 nM Capping Protein in HKEM Buffer and was supplemented with 0.7% BSA, 0.2% methylcellulose, 2.7 mM ATP, 5 mM DTT, 0.2 mM DABCO (motility buffer). When needed the polymerization mix was also supplemented with yeast cofilin (ADF/Cofilin) and/or cyclase associated protein (CAP). The microwells were then closed with mineral oil (Paragon scientific Viscosity Reference Standard RTM13). The flow cell was then sealed with VALAP. Microwells could then be directly imaged under the microscope.

### Imaging

Most of the experiments were done with an epifluorescence system (Ti2 Nikon inverted microscope equipped with a Hamamatsu Orca Flash 4.0 LT Plus Camera or with a Prime BSI Express Camera, Photometrics). Illumination was done with a SOLA III Light Engine (Lumencor). The following objective was used: S Plan Fluor ELWD 20X DIC. Time lapses were acquired with the NIS elements software (version 4.60).

### Image analysis

Images were analyzed with customed macros written in FiJi (Schindelin *et al*, 2012). Data was then processed using customed Python functions and plotted with GraphPad Prism. Mean and standard deviation are represented for all the data. Fits, calculation of correlation coefficient and statistical tests were done with GraphPad Prism. All the statistical information regarding replicates values can be found in figure legends or associated tables.

#### Actin calibration curve

In order to build the calibration curve, we filled microwells of all sizes with a mix of known actin concentration labelled at 5%, with an excess of profilin in order to inhibit polymerization. This gave us “empty” microwells references, from which we could build a calibration curve by plotting the measured intensity of each microwell size against the known number of actin monomers (**Figure S1**). The number of actin monomers was calculated from the volume of each microwell size and the actin concentration.

#### Characterization of structure size and bead velocity

Actin comets were detected with the following procedure. First, threshold was adjusted manually and images were binarized. Actin comet area was obtained from the results of the “Analyze particles” function. Comet area was obtained directly from the result table. Tracking of comets and fluorescent beads was then done with the TrackMate plugin using the thresholding or LoG detectors respectively.

Raw data from Fiji analysis was then analyzed using Python pipelines. For each experiment, essential information such as date, microscope, time interval and protein concentrations were compiled in a reference table. Key parameters such as comet are and bead velocity were compiled for each experiment into tables sorted by individual position, microwell size, number of beads, and timepoint. These tables were finally used to compile analysis between experiment, pooling data by microwell size, timepoint, number of beads, and biochemical conditions. For each analysis, CSV files of interests and the corresponding code were saved in a single folder for traceability.

Finally, Graph Prism was used with the CSV files for proper generating adequate graphs. All further manipulation such as normalization or curve fits were done using Graph Prism available resources.

